# Implication of S-1 biomarkers on aging and prognosis in elderly patients with lung cancer treated by adjuvant S-1 chemotherapy: Accompanying results of the Setouchi Lung Cancer Group Study 1201

**DOI:** 10.1101/2024.11.23.625024

**Authors:** Junichi Soh, Hiromasa Yamamoto, Norihito Okumura, Hiroyuki Suzuki, Masao Nakata, Toshiya Fujiwara, Kenichi Gemba, Isao Sano, Takuji Fujinaga, Masafumi Kataoka, Yasuhiro Terasaki, Nobukazu Fujimoto, Kazuhiko Kataoka, Shinji Kosaka, Motohiro Yamashita, Hidetoshi Inokawa, Masaaki Inoue, Hiroshige Nakamura, Yoshinori Yamashita, Yuta Takahashi, Hidejiro Torigoe, Hiroki Sato, Shuta Tomida, Katsuyuki Hotta, Hiroshige Yoshioka, Satoshi Morita, Keitaro Matsuo, Junichi Sakamoto, Hiroshi Date, Shinichi Toyooka

**Affiliations:** Department of Thoracic Surgery, Okayama University Hospital, 2-5-1 Shikata-cho, Kita-ku, Okayama 700-8558 Japan; Department of Thoracic Surgery, Department of Thoracic Surgery, Osaka Metropolitan University Graduate School of Medicine, 1-4-3 Asahimachi, Abeno-ku, Osaka 545-8585, Japan; Department of Thoracic Surgery, Kurashiki Central Hospital, 1-1-1 Miwa, Kurashiki 710-8602 Japan; Department of Chest Surgery, Fukushima Medical University Hospital, 1 Hikarigaoka, Fukushima 960-1295 Japan; Department of General Thoracic Surgery, Kawasaki Medical School Hospital, 577 Matsushima, Kurashiki 701-0192 Japan; Department of Thoracic Surgery, Hiroshima City Hiroshima Citizens Hospital, 7-33 Motomachi, Naka-ku, Hiroshima 730-8518 Japan; Department of Respiratory Medicine, Chugoku Central Hospital, 148-13 Kamiiwanari, Miyuki-cho, Fukuyama 720-0001 Japan; Department of Respiratory Surgery, Japanese Red Cross Nagasaki Genbaku Hospital, 3-15 Morimachi, Nagasaki 852-8511 Japan; Department of General Thoracic Surgery, National Hospital Organization Nagara Medical Center, 1300-7 Nagara, Gifu 502-8558 Japan; Department of Surgery and Respiratory Center, Okayama Saiseikai General Hospital, 2-25 Kokutai-cho, Kita-ku, Okayama 700-8511 Japan; Department of Respiratory Surgery, Saga Medical Center Koseikan, 400 Nakabaru, Kasemachi, Saga 840-8571 Japan; Department of Medical Oncology and Respiratory Medicine, Okayama Rosai Hospital, 1-10-25 Chikkomidorimachi, Minami-Ku, Okayama 702-8055 Japan; Department of Thoracic Surgery, National Hospital Organization Iwakuni Clinical Center, 1-1-1 Atagomachi, Iwakuni 740-8510 Japan; Department of Thoracic Surgery, Shimane Prefectural Central Hospital, 4-1-1 Himebara, Izumo 693-8555 Japan; Department of Thoracic Surgery, National Hospital Organization Shikoku Cancer Center, 160 Minamiumemotomachi-Ko Matsuyama 791-0280 Japan; Department of Thoracic Surgery, National Hospital Organization Yamaguchi-Ube Medical Center, 685 Higashi-kiwa, Ube 755-0241 Japan; Department of Thoracic Surgery, Shimonoseki City Hospital, 1-13-1 Koyocho, Shimonoseki, 750-8520 Japan; Division of General Thoracic Surgery, Tottori University Hospital, 36-1 Nishi-cho, Yonago 683-8504 Japan; Department of Thoracic Surgery, National Hospital Organization Kure Medical Center and Chugoku Cancer Center, Kure 737-0023 Japan; Center for Comprehensive Genomic Medicine, Okayama University Hospital, Okayama 700-8558 Japan; Center for Innovative Clinical Medicine, Okayama University Hospital, 2-5-1 Shikata-cho, Kita-ku, Okayama 700-8558 Japan; Department of Thoracic Oncology, Kansai Medical University Hospital, 2-3-1 Shinmachi, Hirakata 573-1191 Japan; Department of Biomedical Statistics and Bioinformatics, Kyoto University Graduate School of Medicine, 54 Shogoinkawara-cho, Sakyo-ku, Kyoto 606-8507 Japan; Division of Cancer Epidemiology and Prevention, Aichi Cancer Center Research Institute, 1-1 Kanokoden, Chikusa-ku, Nagoya 464-8681 Japan and Department of Preventive Medicine, Kyushu University Faculty of Medical Sciences, 3-1-1 Maidashi, Higashi-ku, Fukuoka 812-0054 Japan; Tokai Central Hospital, 4-6-2 Sohara Higashijima-cho, Kakamigahara 504-8601 Japan; Department of Thoracic Surgery, Kyoto University Hospital, 54 Shogoinkawara-cho, Sakyo-ku, Kyoto 606-8507 Japan

**Keywords:** non-small cell lung cancer, elderly patients, adjuvant chemotherapy, S-1, EGFR, TP, TS, OPRT, ERCC1, DPD

## Abstract

**BACKGROUNDS:** The management of elderly patients presents several challenges due to age-related declines; however recent recommendations advocate for age not being the sole determinant for adjuvant treatment decisions in patients with non-small cell lung cancer (NSCLC). Aging may alter expression levels of 5-fluorouracil (5-FU) biomarkers.

**METHODS:** Expression changes with aging were explored using The Cancer Genome Atlas (TCGA) database. 5-FU-related biomarker expressions, including thymidylate synthase (TS), dihydropyrimidine dehydrogenase (DPD), orotate phosphoribosyltransferase, epidermal growth factor receptor (EGFR), and excision repair cross-complementation group-1 (ERCC1), were assessed by the quantitative reverse-transcription PCR assays in 89 NSCLCs elderly patients (≥ 75 years old) receiving adjuvant S-1, an oral fluoropyrimidine agent, therapy in a SLCG1201 trial. **RESULTS:** TCGA database analysis (n=955) indicated decreased *TS* expression with aging, particularly in those 75 or older. In the SCLG1201 trial, univariate analysis revealed that high *EGFR* and low *TS* expressions correlated with favorable recurrence-free survival (RFS)(p=0.0264) and overall survival (OS)(p=0.0365), respectively. Multivariable analysis confirmed pathological stage as an independent prognostic factor for both RFS and OS (p=0.0052 and 0.0352, respectively). *EGFR* mutant tumors (n=27) showed significantly higher *DPD* (p=0.0066) and *EGFR* (p<0.0001) expressions, and lower *TS*(p=0.0125) and *ERCC1*(p=0.0015) expressions.

**CONCLUSION:** Despite pathological stage being an independent prognostic factor, high *EGFR* and low *TS* expressions may predict better clinical outcomes in elderly NSCLC patients using adjuvant S-1. The age-related decrease in TS expression supports the potential benefit of 5-FU-based therapy in them compared to younger patients. Further research is warranted to validate these clinical implications.

## INTRODUCTION

Lung cancer is the leading cause of cancer-related death in the world [1]. Radical resection is the standard care for patients with clinical stage I to III non-small cell lung cancer (NSCLC). In Japan, adjuvant chemotherapy is recommended for patients with NSCLC having pathological maximum tumor size with diameter of 2cm or more even with complete tumor resection. In general, elderly individuals are noticed to have poor treatment compliance due to various declines in physical and cognitive functions. The European Organization for Research and Treatment of Cancer and International Society of Geriatric Oncology recommend adjuvant chemotherapy treatment for NSCLC in an elderly population, stating that postoperative adjuvant chemotherapy treatment can achieve favorable survival, and should not be denied to patients based on age [2]. Furthermore, prolonged exposure to some external factors and a decrease in DNA repair functions in elderly individuals may lead to various genetic abnormalities, indicating that molecular biological characteristics implicated in carcinogenesis may differ from those in younger individuals. These aging-specific molecular changes may affect the clinical outcome of systemic therapy in geriatric patients with NSCLC.

S-1 (Taiho Pharmaceutical Co., Ltd, Tokyo, Japan) is an oral fluoropyrimidine agent, consisting of tegafur, a prodrug of 5-fluorouracil (5-FU); gimeracil, a dihydropyrimidine dehydrogenase (DPD) inhibitor which degrades 5-FU; oteracil, a phosphorylation inhibitor in the gastrointestinal tract This products aids in reducing the gastrointestinal toxic effects of 5-FU [3]. In multiple studies, the efficacy of S-1 as monotherapy in elderly patients with advanced NSCLC, and as an alternative treatment of platinum-doublet chemotherapy, has been indicated [4–6]. Furthermore, we recently investigated and reported that the alternate-day and the daily oral administrations of S-1 for 14 consecutive days followed by 7-day rest were feasible in elderly patients with completely resected NSCLC in a phase 2 clinical trial conducted by Setouchi Lung Cancer Group (SLCG1201) [7].

Previous studies have shown that 5-FU sensitivity is influenced by its biomarkers, such as the target molecule, thymidylate synthase (TS) [8]; fluoropyrimidine metabolizing enzymes, such as orotate phosphoribosyl transferase (OPRT) and thymidine phosphatase (TP) [9, 10]; and the 5-FU catabolic enzyme, DPD [11]. Activating mutations of epidermal growth factor receptor (EGFR) gene plays a pivotal role in the treatment of EGFR tyrosine kinase inhibitors (EGFR-TKI). However, previous reports have shown that, by clinical and experimental approach, EGFR mutations had a negative impact on the treatment, using oral uracil-tegafur (UFT: Taiho Pharmaceutical Co., Ltd) [12, 13]. Excision repair cross-complementation group-1 (ERCC1) has been reported to be a crucial factor involved in some DNA repair pathways. Regarding NSCLC, patients who were treated by cisplatin-based chemotherapy, had poor prognosis with the expression of ERCC1; low ERCC1 expression correlated with better prognosis in patients with gastric and colon cancer following treatment by 5-FU [14]. However, there has not been much research done on the relationship between the expression level of ERCC1 and the therapeutic effect of 5-FU chemotherapy in patients with NSCLC.

In this study, we aim to investigate the aging effect on the expression level of 5-FU biomarkers using a public database, and to conduct a biomarker study in predicting clinical outcomes of adjuvant chemotherapy with S-1 oral administration in elderly patients with NSCLC who underwent complete resection in the SCLG1201 trial.

## MATERIALS AND METHODS

### Analysis of The Cancer Genome Atlas (TCGA) datasets

RNA-Seq gene expression data for TCGA samples were collected from the Genomic Data Commons Portal (https://portal.gdc.cancer.gov). We downloaded two datasets named TCGA-LUAD and TCGA-LUSC. The z-scores of each gene (*DPYD*, *TYMP*, *TYMS*, *UMPS*, *ERCC1*, and *EGFR*), *EGFR* mutational status, and clinicopathological data such as age, sex, histology, and race were retrieved.

### Patients’ selection, ethics approval and consent to participate

We enrolled patients who had agreed to participate from May 2012 to April 2016 in both the SLCG1201 (University hospital Medical Information Network ID: UMIN000007819) [7] and its subsidiary study. Patients provided written informed consents for both the main and accompanying SLCG1201 studies. The SCLG1201 study was approved by Okayama University Certified Review Board (approval number: CRB18-011), followed by the confirmation of each participating institution. The accompanying study of SCLG1201 was approved by the Ethics Committee, Okayama University Graduate School of Medicine, Dentistry and Pharmaceutical Sciences and Okayama University Hospital, Okayama, Japan (approval number: Rin1945). It was also approved by the institutional review board of each participating institution. The study’s data was managed by the SLCG1201 data center at a non-profit organization; the Epidemiological and Clinical Research Information Network (ECRIN), Kyoto, Japan.

### Sample collection and preparation

Surgical specimens were fixed in 10% formaldehyde and embedded in paraffin. The resulting formaldehyde-fixed and paraffin-embedded (FFPE) blocks, including the primary tumor, were sectioned and overlaid on non-coating slides. Two slides received the five μm-thick sections, and five, the ten μm-thick sections, at each participating institute. These slides were immediately sent to the Department of Thoracic Surgery, Okayama University Hospital. After confirmation of the quality of each section and anonymization, these slides were immediately sent to FALCO biosystems Ltd. (Kyoto, Japan).

Representative hematoxylin and eosin-stained slides were made from the FFPE slide; five μm-thick, and reviewed by a pathologist for a manual macrodissection of the tumor tissue. Tumor tissue was selected from the FFPE slides which were ten μm-thick, and dissected using a scalpel. RNA was isolated from dissected tumor tissue using RNeasy FFPE Kit (Qiagen, Chatsworth, GA). cDNA was prepared using High-Capacity Reverse Transcription Kit (Life Technologies, Foster City, CA), according to the manufacturer’s instruction.

### Quantitative reverse-transcription PCR

Following TaqMan assay-based pre-amplification, the expression levels of six genes, which are: *TP*, *TS*, *DPD*, *OPRT*, *ERCC1*, and *EGFR*, were determined using TaqMan real-time PCR (TaqMan array card; Life Technologies). Briefly, 2.5 µl of cDNA was pre-amplified using TaqMan PreAmp Master Mix (2×) (Life Technologies) and a pool of TaqMan® Gene Expression Assays (0.2×) in a 10-µL PCR reaction. The pre-amplification cycling conditions were as follows: 95°C for 10 min followed by 14 cycles of 95°C and 60°C for 15 s and 4 min, respectively. An amplified cDNA sample was diluted 20 times in TE buffer. Amplified cDNA (25 µL) was added to 25 µL RNase-free water and 50 µL of 2× TaqMan Gene Expression Master Mix (Life Technologies). The mixture was transferred to a loading port for the TaqMan array card (LDA). The array card was centrifuged twice, sealed, and PCR amplification was performed using the Applied Biosystems Prism 7900HT Sequence Detection System (Life Technologies) under the following thermal cycling conditions: 50°C and 94.5°C for 2 and 10 min, respectively, followed by 40 cycles of 97°C and 59.7°C for 30 s and 1 min, respectively. The array card included ACTB as a reference based on its proven role as the housekeeping gene. The assay IDs used in the array card are shown in Supplemental table 1. The cycle threshold (Ct) value, which is inversely proportional to the amount of cDNA, was calculated. The relative mRNA expression levels were stated as the ratios (the differences between the Ct values) between the gene of interest and the reference gene.

### Statistical analysis

Significant differences between the categorized groups were compared using the Chi-square or Fisher’s exact test and for the analyses of continuous variables, the Mann-Whitney U test was used. We obtained the follow-up data from the original study of SLCG1201 [7]. Univariate and multivariate analyses of overall and recurrence-free survival (OS and RFS) were performed using the Kaplan-Meier method with log-rank and the Cox proportional hazard testing. We defined p < 0.05 as the threshold for statistical significance. All statistical analyses were performed using the JMP 9.0.2 Program for Windows (SAS Institute Inc., Cary, NC, USA) and GraphPad Prism 9 (GraphPad Software, La Jolla, CA, USA).

## RESULTS

### Impact of aging on the expression of six genes in the public database

We investigated the impact of aging on the expression of six genes using the TCGA database. RNA-Seq gene expression data were obtained from 955 samples on the TCGA, comprising 776 samples from patients under 75 years, and 179 samples from patients over 75 years (Supplemental Table 2). The expression levels of the six genes were compared across age (Figure 1). *TS* expression levels significantly decreased with age, especially in patients 75 years or older (p = 0.0155). In contrast, *DPD* and *TP* expression levels increased with age, but the expression levels of these two genes between those under 75 and 75 years or older, were not statistically significant.

**Fig. 1.**
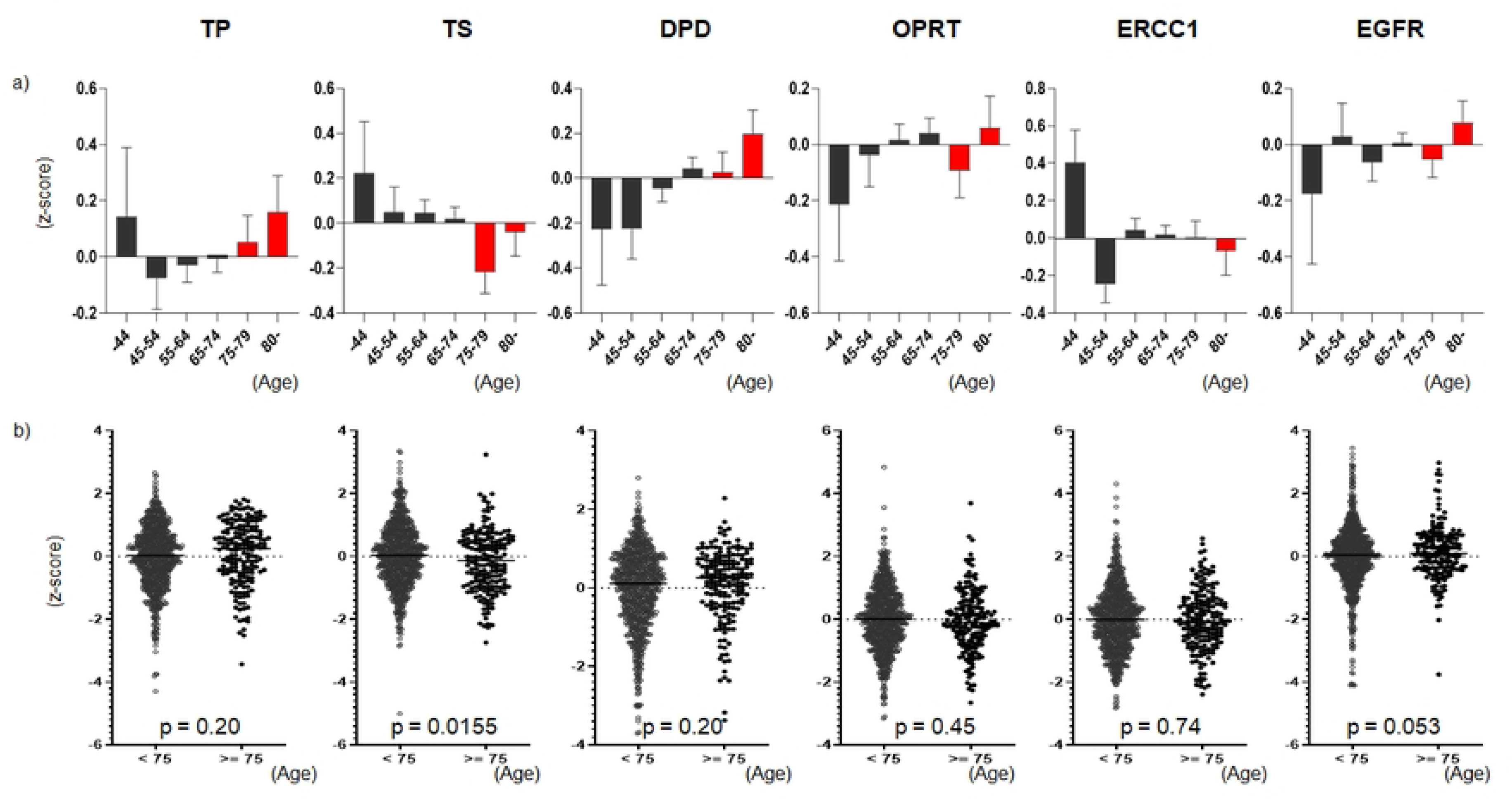
Impact of aging on mRNA expression levels of six genes. We downloaded two datasets named TCGA-LUAD and TCGA-LUSC from the Genomic Data Commons Data Portal (https://portal.gdc.cancer.gov). a) the mean and standard deviation of the z-scores for the six genes are shown by age group based on 10-year increments. b) data from all 955 samples were plotted and compared between patients of less than 75 years old (n = 776) and those of 75 years old or more (n = 179). The Mann-Whitney U test was used for comparison between the two groups.

### Patient characteristics

Of the 97 patients who received the allocated intervention, 90 agreed to participate in the accompanying SLCG1201 study. However, one sample was excluded from the study because of the lack of tumors in the provided slides, leaving 89 patients enrolled in the study. The baseline characteristics of the enrolled patients are summarized in Table 1. The median age was 77 years old (range, 75 - 87), and 65 (73.0%) were men. Fifty-seven (64.0%) patients had adenocarcinoma histology and 54 (60.7%) were pathological stage (pStage) IA (T1bN0M0) / IB. *EGFR* mutation was present in 27 (30.3%) patients.

**Table 1.**
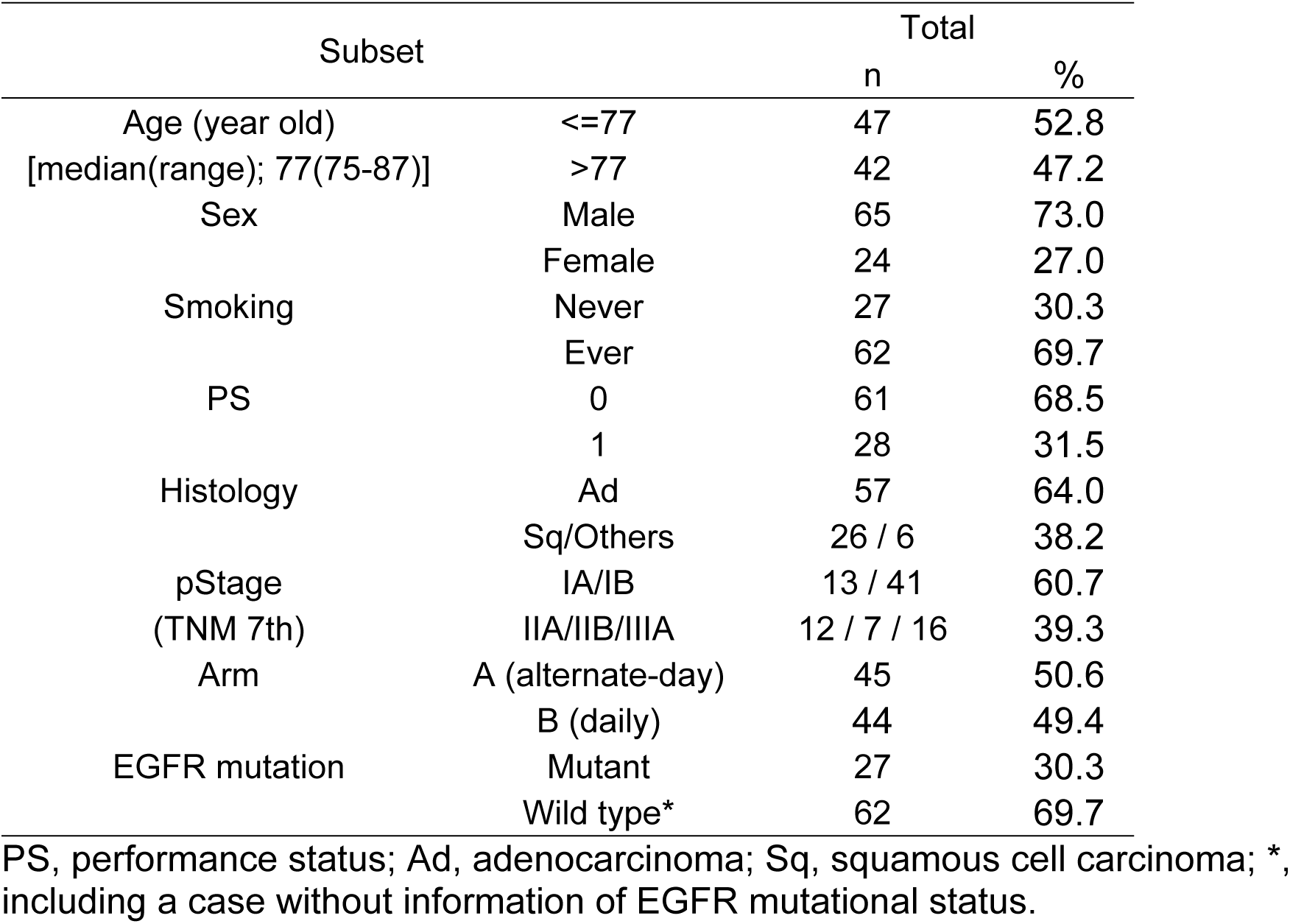

### Relative expression levels of six genes and EGFR mutations

The results of the relative mRNA expression levels of all genes were shown in Figure 2. The median Ct value of *ACTB* of all 89 samples was 15.69 (range, 11.65 to 26.66), indicating that the mRNA quality of all samples was adequate. However, three samples could not be amplified in the specific genes: one sample for *TP* and *EGFR* (the Ct value of *ACTB* in this sample, 26.66), and two for *OPRT* (the Ct value of ACTB,21.72 and 19.88, respectively). The correlation between the relative mRNA expression levels of six genes were investigated to find that there was no significant correlation between them (Supplemental figure 1).

**Fig. 2.**
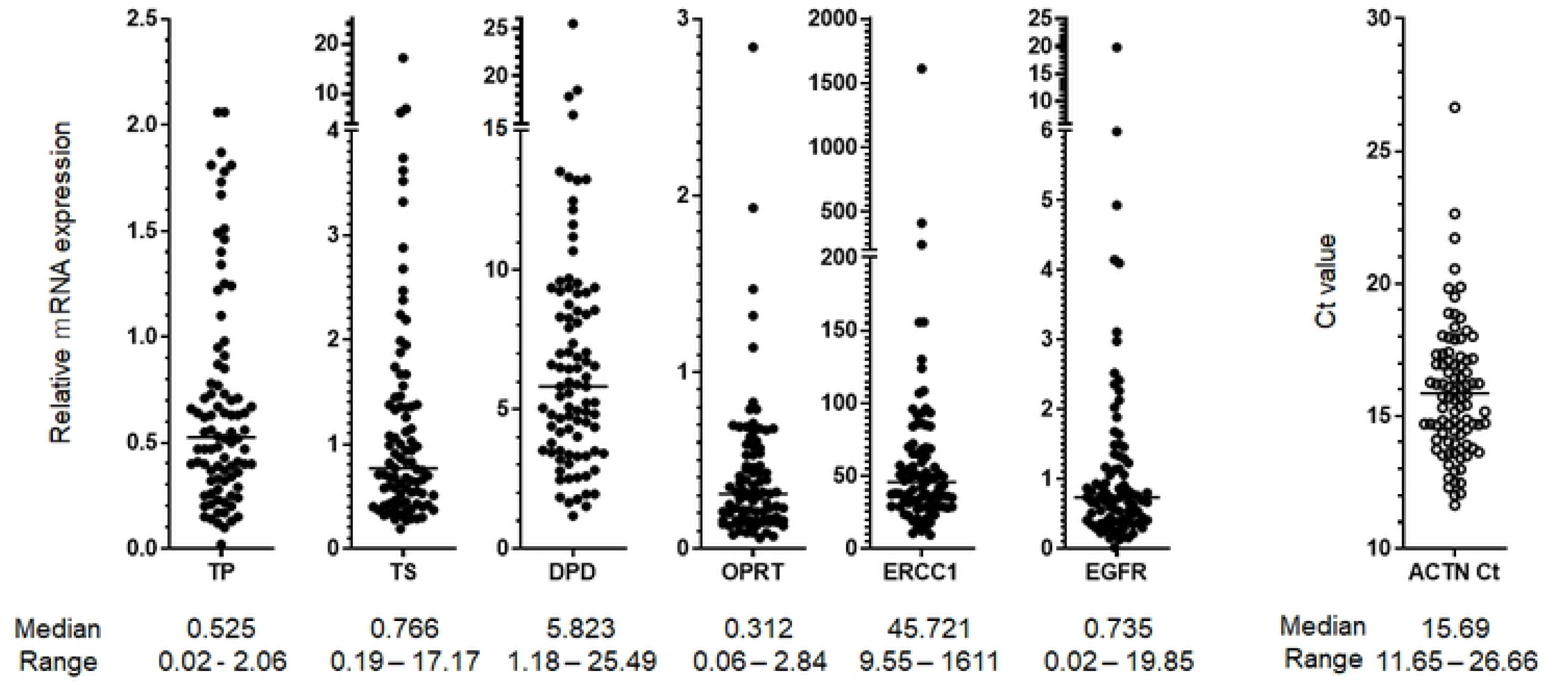
Relative mRNA expression of six genes. The relative mRNA expression of each six gene was plotted for each sample. The Ct value of *ACTB* (the housekeeping gene) was also plotted as a reference.

The relationships between clinicopathological factors and molecular profiles were indicated in Table 2. *EGFR* mutations were significantly frequent in females, never smokers, patients with poor performance status (PS), and patients with adenocarcinoma, as previously reported [15–17]. The relative mRNA expressions of *TS* and *OPRT* were significantly higher in males; contrary, that of *DPD* and *EGFR* were significantly higher in females. The relative mRNA expression of *TS* was significantly higher in ever smokers; however, that of *DPD* and *EGFR* were significantly higher in never smokers. The relative mRNA expressions of *TS, OPRT* and *ERCC1* were significantly higher in patients with non-adenocarcinoma histology; in contrast, that of *DPD* was significantly higher in patients with adenocarcinoma histology.

**Table 2.** Relationship between clinicopathological factors and each molecular profile.

| Subsets |  | EGFR mutation |  |  | TP |  | TS |  | DPD |  | OPRT |  | ERCC1 |  | EGFR |  |
| --- | --- | --- | --- | --- | --- | --- | --- | --- | --- | --- | --- | --- | --- | --- | --- | --- |
|  |  | Mutant |  |  | median | p | median | p | median | p | median | p | median | p | median | p |
|  |  | n | % | p |  |  |  |  |  |  |  |  |  |  |  |  |
| Age<br>(year old) | ≤77 | 17 | 36.2 | 0.25 | 0.558 | 0.4496 | 0.825 | 0.6104 | 6.155 | 0.3121 | 0.323 | 0.9864 | 46.746 | 0.4545 | 0.772 | 0.2102 |
|  | >77 | 10 | 23.8 |  | 0.472 |  | 0.703 |  | 5.700 |  | 0.279 |  | 43.152 |  | 0.684 |  |
| Sex | Male | 12 | 18.5 | **0.0002** | 0.548 | 0.4258 | 0.987 | **0.0016** | 5.226 | **0.0109** | 0.346 | **0.0204** | 49.652 | 0.0581 | 0.671 | **0.044** |
|  | Female | 15 | 62.5 |  | 0.470 |  | 0.500 |  | 8.219 |  | 0.232 |  | 36.749 |  | 0.988 |  |
| Smoking | Never | 14 | 51.9 | **0.0056** | 0.559 | 0.8281 | 0.532 | **0.0047** | 8.116 | **0.0024** | 0.250 | 0.126 | 37.629 | 0.0908 | 1.062 | **0.0346** |
|  | Ever | 13 | 21.0 |  | 0.502 |  | 0.982 |  | 5.153 |  | 0.322 |  | 50.381 |  | 0.669 |  |
| PS | 0 | 14 | 23.0 | **0.045** | 0.447 | **0.0172** | 0.825 | 0.2295 | 5.892 | 0.5598 | 0.255 | **0.0074** | 49.652 | **0.0157** | 0.684 | 0.0921 |
|  | 1 | 15 | 53.6 |  | 0.667 |  | 0.685 |  | 5.406 |  | 0.466 |  | 33.552 |  | 0.850 |  |
| Histology<br>(Registered) | Ad | 26 | 45.6 | **<0.0001** | 0.546 | 0.2214 | 0.643 | **0.0003** | 6.560 | **0.0044** | 0.235 | **0.0175** | 37.551 | **0.0003** | 0.742 | 0.6403 |
|  | Non-Ad | 1 | 3.1 |  | 0.472 |  | 1.239 |  | 3.857 |  | 0.405 |  | 61.305 |  | 0.675 |  |
| pStage<br>(Registered) | IA/IB | 19 | 35.2 | 0.25 | 0.629 | 0.138 | 0.702 | 0.1202 | 6.586 | **0.0089** | 0.318 | 0.7997 | 41.738 | 0.1059 | 0.797 | 0.1766 |
|  | IIA/IIB/IIIA | 8 | 22.9 |  | 0.468 |  | 0.940 |  | 4.610 |  | 0.268 |  | 49.241 |  | 0.643 |  |
| Arm | A | 23 | 51.1 | 1 | 0.522 | 0.97 | 0.754 | 0.9346 | 5.823 | 0.8761 | 0.315 | 0.6527 | 45.721 | 0.964 | 0.730 | 0.8314 |
|  | B | 21 | 48.8 |  | 0.557 |  | 0.850 |  | 5.788 |  | 0.298 |  | 45.151 |  | 0.742 |  |
| EGFR<br>mutation | Mutant | - | - | - | 0.528 | 0.86 | 0.583 | **0.0125** | 8.116 | **0.0066** | 0.245 | 0.31 | 35.13 | **0.0015** | 1.148 | **<.0001** |
|  | Wild type* | - | - |  | 0.522 |  | 0.982 |  | 5.066 |  | 0.323 |  | 50.38 |  | 0.566 |  |

PS, performance status; Ad, adenocarcinoma; Arm A is alternative day administration of S-1; Arm B is two-week daily administration of S-1; \*, including a case without information of EGFR mutational status.

*EGFR* mutations are important driver mutations in NSCLC, hence, we investigated the interrelationship between mRNA expression levels of six genes and *EGFR* mutations. In our study, *EGFR* mutations were significantly associated with higher relative mRNA expressions of *DPD* (p = 0.0066) and *EGFR* (p < 0.0001), and lower relative mRNA expressions of *TS* (p = 0.0125) and *ERCC1* (p = 0.0015) (Table 2).

### Prognostic impact of clinicopathological factors and molecular profiles

Using the cut-off values of the median level of relative mRNA expressions of six genes, we divided the patients into two groups for further analysis.

Regarding the univariate analysis of RFS, never smokers, patients with pStage I, and patients harboring higher relative *EGFR* mRNA expression showed significantly better RFS than the others (Figure 3, and Table 3a). Multivariate analysis including the factors with a p-value of less than 1 showed that pStage I was an independent favorable prognostic factor for RFS (Table 3a).

**Fig. 3.**
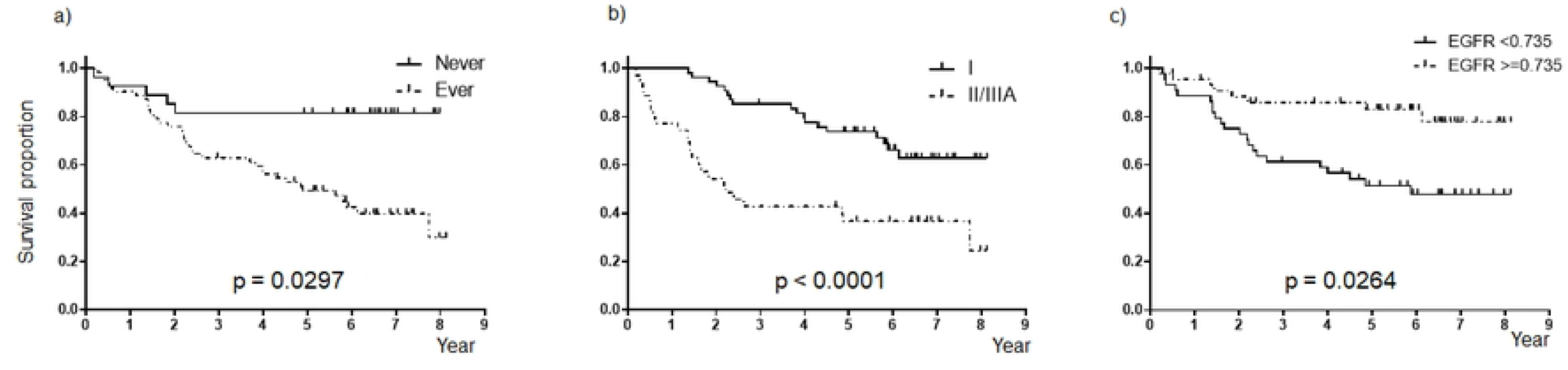
Recurrence-free survival. a) smoking status, b) pathological stage, c) relative mRNA expression of *EGFR*.

**Table 3.** Prognosis.

a) Recurrence-free survival
| Subsets |  | Univariate |  |  | Multivariate |  |  |  |
| --- | --- | --- | --- | --- | --- | --- | --- | --- |
|  |  | HR | p | 95%CI | HR | p | 95%CI |  |
| Age (year old) | (>77 vs. ≤77) | 0.99 | 0.97 | 0.47 to 2.02 | - | - | - | - |
| Sex | (Male vs. Female) | 2.20 | **0.081** | 0.91 to 6.54 | 1.09 | 0.90 | 0.28 to 4.84 |  |
| Smoking | (Ever vs. Never) | 2.64 | **0.0297** | 1.09 to 7.84 | 3.18 | 0.09 | 0.85 to 14.13 |  |
| PS | (1 vs. 0) | 1.13 | 0.75 | 0.51 to 2.36 | - | - | - | - |
| Histology | (non-Ad vs. Ad) | 1.83 | 0.10 | 0.88 to 3.76 | - | - | - | - |
| pStage | (II/IIIA vs. I) | 4.62 | **<.0001** | 2.21 to 10.32 | 2.51 | **0.0052** | 1.32 to 4.85 |  |
| EGFR mutation | (Wild type* vs. Mutant) | 1.25 | 0.58 | 0.58 to 3.00 | - | - | - | - |
| TP expression* | (<0.525 vs. ≥0.525) | 0.70 | 0.25 | 0.37 to 1.29 | - | - | - | - |
| TS expression | (<0.766 vs. ≥0.766) | 0.77 | 0.41 | 0.41 to 1.43 | - | - | - | - |
| DPD expression | (<5.825 vs. ≥5.825) | 1.58 | 0.15 | 0.85 to 3.02 | - | - | - | - |
| OPRT expression** | (<0.312 vs. ≥0.312) | 1.04 | 0.89 | 0.56 to 1.94 | - | - | - | - |
| ERCC1 expression | (<45.721 vs. ≥45.721) | 0.99 | 0.96 | 0.53 to 1.83 | - | - | - | - |
| EGFR expression* | (<0.735 vs. ≥0.735) | 2.03 | **0.0264** | 1.09 to 3.92 | 1.25 | 0.51 | 0.64 to 2.54 |  |
| Arm | (A vs. B) | 1.25 | 0.54 | 0.61 to 2.59 | - | - | - | - |

b) Overall survival
| Subsets |  | Univariate |  |  | Multivariate |  |  |  |
| --- | --- | --- | --- | --- | --- | --- | --- | --- |
|  |  | HR | p | 95%CI | HR | p | 95%CI |  |
| Age (year old) | (>77 vs. ≤77) | 0.98 | 0.947 | 0.47 to 2.01 | - | - | - | - |
| Sex | (Male vs. Female) | 13.55 | **<.0001** | 2.90 to 241.39 | 2.91 | 0.41 | 0.28 to 77.86 |  |
| Smoking | (Ever vs. Never) | 16.00 | **<.0001** | 3.42 to 285.16 | 6.80 | 0.09 | 0.80 to 177.85 |  |
| PS | (1 vs. 0) | 1.19 | 0.66 | 0.53 to 2.49 | - | - | - | - |
| Histology | (non-Ad vs. Ad) | 2.44 | **0.0171** | 1.18 to 5.06 | 1.63 | 0.28 | 0.68 to 4.15 |  |
| pStage | (II/IIIA vs. I) | 2.46 | **0.0143** | 1.20 to 5.17 | 2.26 | **0.0352** | 1.06 to 4.96 |  |
| EGFR mutation | (Wild type* vs. Mutant) | 2.40 | **0.0509** | 1.00 to 7.11 | 0.72 | 0.58 | 0.24 to 2.50 |  |
| TP expression* | (<0.525 vs. ≥0.525) | 0.80 | 0.53 | 0.38 to 1.63 | - | - | - | - |
| TS expression | (<0.766 vs. ≥0.766) | 0.46 | **0.0365** | 0.21 to 0.95 | 0.78 | 0.52 | 0.35 to 1.66 |  |
| DPD expression | (<5.825 vs. ≥5.825) | 1.88 | **0.089** | 0.91 to 4.10 | 0.86 | 0.73 | 0.35 to 2.10 |  |
| OPRT expression** | (<0.312 vs. ≥0.312) | 0.83 | 0.61 | 0.40 to 1.70 | - | - | - | - |
| ERCC1 expression | (<45.721 vs. ≥45.721) | 0.61 | 0.18 | 0.28 to 1.25 | - | - | - | - |
| EGFR expression* | (<0.735 vs. ≥0.735) | 1.73 | 0.14 | 0.83 to 3.76 | - | - | - | - |
| Arm | (A vs. B) | 1.57 | 0.22 | 0.77 to 3.36 | - | - | - | - |

PS, performance status; AD, adenocarcinoma; Arm A is alternative day administration of S-1; Arm B is two-week daily administration of S-1; HR, hazard ratio; CI, confidential interval; \*, including a case without information; \*\*, including two cases without information

In univariate analysis of OS; females, never smokers, adenocarcinoma histology, pStage I, and those harboring lower relative *TS* mRNA expression showed significantly better OS than the others (Figure 4 and Table 3b). Multivariate analysis including the factors with p-value ≤ 1 showed that pStage I was an independent favorable prognostic factor for overall survival (Table 3b).

**Fig. 4.**
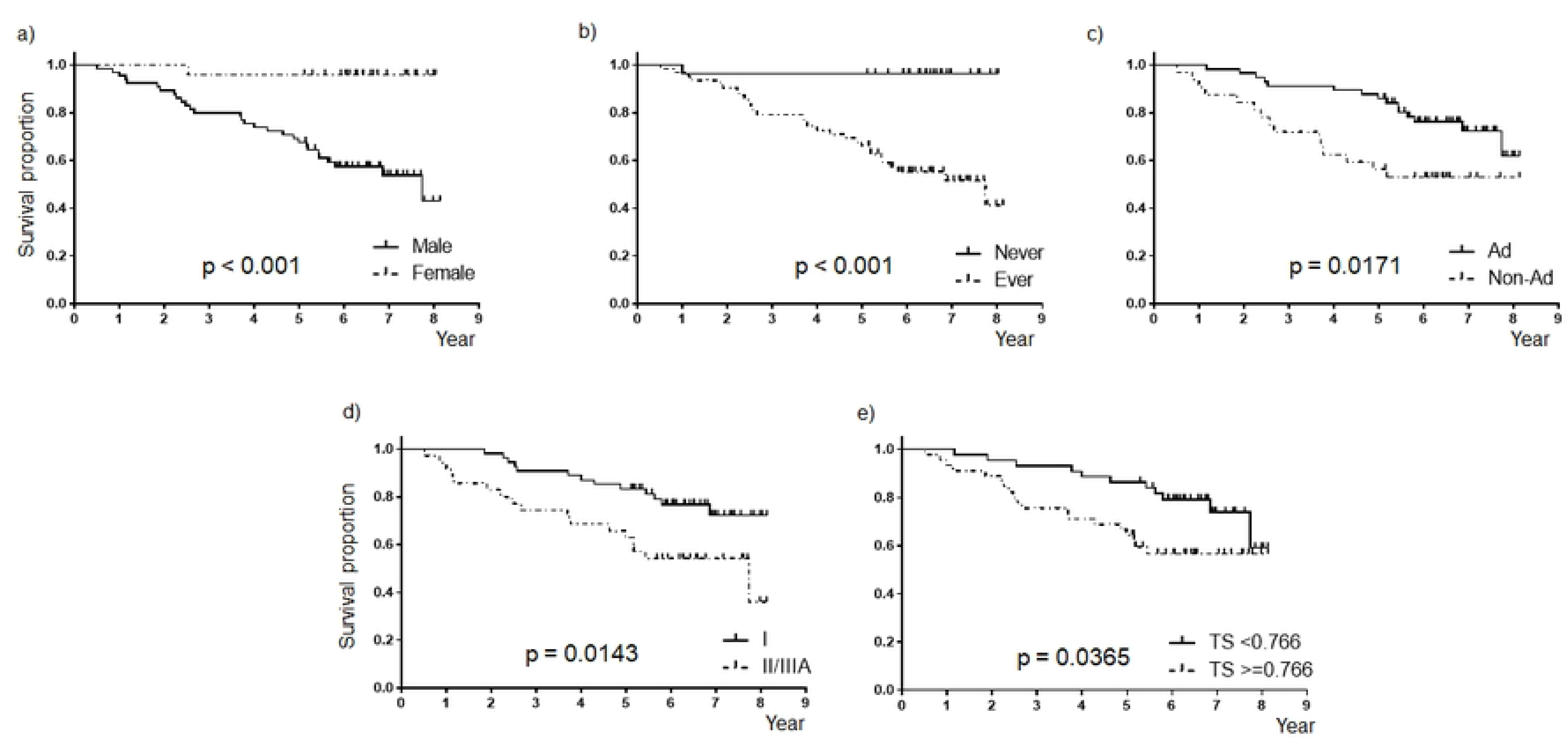
Overall survival. a) sex, b) smoking status, c) histology, d) pathological stage, e) relative mRNA expression of *TS*.

## DISCUSSION

Aging affects gene expression by acting on the splicing of the mRNA or on genetic regulation [18, 19]. Thousands of genes have been identified to be associated with age in multiple tissues [20, 21]. To our knowledge, the six genes investigated in our study have not been linked with aging. However, RNA-Seq gene expression data of the TCGA database indicated that *TS* expression level may be changed according to aging. These findings suggest that expression levels of 5-FU biomarkers also change with age, potentially impacting the clinical outcome of adjuvant treatment in elderly patients with NSCLC compared with younger patients.

This study examined the mRNA expression of six 5-FU-related markers considered as treatment biomarkers in S-1 oral therapy for pStage IA (2 cm) to IIIA in patients aged 75 years or older. We investigated their association with clinicopathological factors and prognosis. Notably, high expression of *EGFR* and *DPD* genes and low expression of *ERCC1* and *TS* genes were significantly associated with the presence of *EGFR* mutation. Univariate analysis for RFS and OS revealed low *EGFR* expression was as an unfavorable factor for RFS, and high *TS* expression was an unfavorable factor for OS. In multivariate analysis, pStage II/III was identified as an independent unfavorable factor for both RFS and OS.

Our study revealed that *EGFR* mutations were significantly associated with higher relative mRNA expressions of *EGFR* and *DPD* genes and lower relative mRNA expressions of *ERCC1* and *TS* genes. Consistent with our findings related to *EGFR*. Previous studies have shown that mutant alleles specific amplification which induces high mRNA expression of *EGFR* was frequently observed in the patients harboring *EGFR* mutations [22]. Thereby resulting in the significant association between the presence of mutation and the high mRNA expression of *EGFR* gene which has been also indicated by other previous reports [23, 24]. Regarding *DPD*, previous studies have shown that high expression was correlated with *EGFR* mutation in both clinical samples and on the cell lines [25]. Also, the *EGFR* signal cascade regulated *DPD* expression via Sp1 in *EGFR* mutant cells [26]. Regarding *ERCC1*, an experimental study showed that increased DNA damage and reduced damage repair (ERCC1 and RAD51 foci formation) was observed in *EGFR* exon 19 deletion mutation cells, compared with *EGFR* wild-type cells [27]. Furthermore, *ERCC1* expression was increased by inhibition of *EGFR* exon 19 deletion signals; on the other hand, *ERCC1* expression was decreased by blockage of wild-type EGFR signals. Multiple studies have indicated that NSCLC specimens with *EGFR* activating mutations tend to have low *ERCC1* mRNA levels [28–31]. ERCC1 and TS are essential for DNA synthesis and repair, and *TS* expression was decreased in *EGFR* mutant lung cancer specimens as presented in our study [28, 31].

Univariate analysis found that low *EGFR* mRNA expression and high *TS* mRNA expression were significantly associated with unfavorable RFS and OS, respectively. To our knowledge, the prognostic impact of *EGFR* mRNA expression has not been well-investigated in patients who received 5-FU adjuvant chemotherapy after complete resection. However, we have previously reported that adjuvant UFT can significantly improve the prognosis in patients without *EGFR* mutation but not in patients with *EGFR* mutation along with experimental confirmation [12]. These findings were also suggested by a recent similar study [13]. In contrast, a meta-analysis of 2972 patients from 18 studies, conducted before EGFR-tyrosine kinase inhibitors were developed, showed that overexpression of EGFR was not associated with favorable prognosis (combined hazard ratio of 1.14 with 95% confidence interval of 0.97 to1.34; p = 0.103) [32]. Several studies and meta-analysis indicated that low TS protein expression was significantly associated with favorable prognosis in lung cancer patients who received S-1-related [33] and pemetrexed-related [34] therapies. Furthermore, TS overexpression was associated with poor prognosis of many cancer patients treated with 5-FU and UFT, including NSCLC [33, 35, 36] and gastrointestinal cancers [37].

There are some limitations found in this study. Although the number of patients enrolled in this study is limited, all enrolled patients were monitored until death or for at least 5 years from the registration. mRNA detection was performed using a commercial-based real-time PCR assay using primers for a specific lesion of the target gene. However, we did not confirm the correlation between mRNA expression level and protein expression level. As a preliminary study, we confirmed the correlation of mRNA and protein expression levels for a part of target genes such as *TP*, *DPD*, and *EGFR* (data not shown). Experimental and clinical studies have identified that a non-synonymous SNP 538G>A f MRP8/ABCC11, one of the ABC transporters for which 5-FU, methotrexate (MTX), and pemetrexed (PEM) are substrates, as a potential biomarker for S-1 treatment [38, 39]. Future studies will clarify the clinical implications of these candidate biomarkers.

In conclusion, *TS* expression was significantly decreased with aging in the public database. Univariate analysis showed low *TS* and high *EGFR* expression levels demonstrated as better prognostic markers among 5-FU-related molecules in elderly patients with NSCLC who underwent radical resection followed by adjuvant therapy with oral administration of S-1. Although pStage was identified as an independent prognostic factor through multivariable analysis. These findings suggested that elderly patients with NSCLC exhibited low *TS* expression, that may have resulted in a better clinical outcome with 5-FU-based treatment, including S-1 in this population.

## ACKNOWLEDGEMENTS

We thank Ms. Yumi Miyashita [a non-profit organization Epidemiological & Clinical Research Information Network (ECRIN)] for data management. We would like to thank Editage (www.editage.jp) for English language editing.

## FUNDING

This research did not receive any specific grant from funding agencies in the public, commercial, or not-for-profit sectors.

## AVAILABILITY OF DATA AND MATERIALS

Data and materials of this work are available from the corresponding author upon request.

## Abbreviations

NSCLC: non-small cell lung cancer;
CI: confidence interval;
SD: standard deviation;
EGFR: epidermal growth factor receptor;
DPD: dihydropyrimidine dehydrogenase;
DPYD: the gene symbol of DPD;
TP: thymidine phosphorylase;
TYMP: the gene symbol of TP;
TS: thymidylate synthetase;
TYMS: the gene symbol of TS;
OPRT: orotate phosphoribosyl transferase;
UMPS: the gene symbol of OPRT;
ERCC1: excision repair cross-complementing 1 (the gene symbol as well);
ACTB: the gene symbol of actin beta;
RFS: relapse-free survival;
OS: overall survival

## Supplementary Figure Legends

**Supplementary Fig. 1.** Correlation between relative mRNA expression levels of six genes.

## Notes

### Competing Interest Statement

I have read the journal's policy and the authors of this manuscript have the following potential competing interests out of this work: Junichi Soh reports a relationship with Chugai Pharmaceutical Co Ltd, Johnson & Johnson MedTech, Medtronic Inc, Intuitive Surgical Inc, CSL Behring LLC, and Olympus Corporation that includes: funding grants and speaking and lecture fees. Hiromasa Yamamoto reports a relationship with Chugai Pharmaceutical Co Ltd that includes: funding grants. Nobukazu Fujimoto reports a relationship with Merck & Co Inc, Ono Pharmaceutical Co Ltd, Nippon Kayaku Co Ltd, Chugai Pharmaceutical Co Ltd, Taiho Pharmaceutical Co Ltd, and Nippon Boehringer Ingelheim Co Ltd that includes: speaking and lecture fees. Katsuyuki Hotta reports a relationship with AstraZeneca Pharmaceuticals LP, Chugai Pharmaceutical Co Ltd, Eli Lilly Japan KK, MSD, Bristol-Myers Squibb Company, Ono Pharmaceutical Co Ltd, Boehringer Ingelheim Pharmaceuticals Inc, Nippon Kayaku Co Ltd, Amgen Inc, Taiho Pharmaceutical Co Ltd, Merck & Co Inc, and AbbVie Inc that includes: consulting or advisory, funding grants, and speaking and lecture fees. Hiroshige Yoshioka reports a relationship with Amgen Inc, AstraZeneca, Bristol Myers Squibb Co, Chugai Pharmaceutical Co Ltd, Daiichi Sankyo Inc, Delta-Fly Pharma, Eli Lilly and Company, Kyowa Kirin Co Ltd, Merch Biopharma, MSD, Boehringer Ingelheim Co Ltd, Nippon Kayaku Co Ltd, Nipro Pharma Corporation, Novartis Pharmaceuticals, Ono Pharmaceutical Co Ltd, Otsuka Pharmaceutical Co Ltd, Pfizer, Taiho Pharmaceutical Co Ltd, and Takeda Pharmaceutical Company Limited that includes: consulting or advisory, and speaking and lecture fees. Hiroshi Date reports a relationship with Johnson and Johnson Ltd, AstraZeneca Pharmaceuticals LP, and TriMed Inc that includes: speaking and lecture fees and funding grants. Shinichi Toyooka reports a relationship with Chugai Pharma, Daiichi Sankyo, Astellas Pharma, Boehringer Ingelheim, Taiho Pharmaceutical, Ono Pharmaceutical, Bayer, Medtronic, Johnson & Johnson, Kyorin, Eli Lilly, Novartis, AstraZeneca, Merck & Co, Guardant Health, and Eli Lilly that includes: consulting or advisory, funding grants, and speaking and lecture fees.

